# Limit distribution of the quartet balance index for Aldous’s *β* ≥ 0-model

**DOI:** 10.1101/277376

**Authors:** Krzysztof Bartoszek

## Abstract

This paper builds up on T. Martínez-Coronado, A. Mir, F. Rossello and G. Valiente’s work “A balance index for phylogenetic trees based on quartets”, introducing a new balance index for trees. We show here that this balance index, in the case of Aldous’s *β* ≥ 0-model, convergences weakly to a distribution that can be characterized as the fixed point of a contraction operator on a class of distributions.

## 1 Introduction

Phylogenetic trees are key to evolutionary biology. However, they are not easy to summarize or compare as it might not be obvious how to tackle their topologies, understood as the internal branching structure of the trees. Therefore, many summary indices have been proposed in order to “project” a tree into ℝ. Such indices have as their aim to quantify some property of the tree and one of the most studied properties is the symmetry of the tree. Tree symmetry is commonly captured by a balance index. Multiple balance indices have been proposed, Sackin’s (Sackin, 1972), Colless’ (Colless, 1982) or the total cophenetic index (Mir et al., 2013). A compact introduction to phylogenetics, containing in particular a list of tree asymmetry measures (p. 562-564), can be found in Felsenstein (2004)’s book. This work accompanies a newly proposed balance index—the quartet index (QI, Martínez-Coronado et al., 2018b).

One of the reasons for introducing summary indices for trees is to use them for significance testing—does a tree come from a given probabilistic model. Obtaining the distribution (for a given *n*-number of contemporary species, i.e. leaves of the tree, or in the limit *n* → ∞) of indices is usually difficult and often is done only for the “simplest” Yule (pure–birth Yule, 1924) tree case and sometimes uniform model (see e.g. Aldous, 1991; Steel and McKenzie, 2001).

Using the contraction method, central limit theorems were found for various balance indices, like the total cophenetic index (Yule model case Bartoszek, 2018) and jointly for Sackin’s and Colless’ (in the Yule and uniform model cases Blum et al., 2006). Furthermore, Blum and François (2006) showed that Sackin’s index has the same weak limit as the number of comparisons of the quicksort algorithm (Hoare, 1962), both after normalization of course.

Chang and Fuchs (2010) consider the number of occurrences of patterns in a tree, where a pattern is understood as “any subset of the set of phylogenetic trees of fixed size *jk*. For a tree with *n* leaves such a pattern will satisfy the recursion

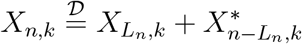

where *X_n,k_*, 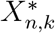 and *L_n_* are independent, 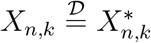 and *L_n_* is the size of the left subtree branching from the root. For the Yule and uniform models they derived central limit theorems (normal limit distribution) with Berry–Esseen bounds and Poisson approximations in the total variation distance.

Even though the pure–birth model seems to be very widespread in the phylogenetics community, more complex models need to be studied, especially in the context of tree balance. From Roch and Snir (2013)’s Lemma 4 it can be deduced that Yule trees have to be rather balanced— as the maximum quartet weight (maximum of number of randomly placed marks along branches over induced subtrees on four leaves) is asymptotically proportional to the expectation of the tree’s height.

In this work here, using the contraction method, we show convergence in law of the (scaled and centred) quartet index and derive a representation (as a fixed point of a particular contraction operator) of the weak–limit. Remarkably, this is possible not only for the Yule tree case but for Aldous’s more general *β*–model (in the *β* ≥ 0 regime).

The paper is organized as follows. In Section 2 we introduce Aldous’s *β*–model and the quartet index. In Section 3 we prove our main result— Thm. 3.1 via the contraction method. When studying the limit behaviour of recursive–type indices for pure–birth binary trees one has that for each internal node the leaves inside its clade are uniformly split into to sub–clades as the node splits. However, in Aldous’s *β*–model this is not the case, the split is according to a BetaBinomial distribution, and a much finer analysis is required to show weak–convergence, with *n*, of the recursive–type index to the fixed point of the appropriate contraction. Theorem 3.1 is not specific for the quartet index but covers a more general class of models, where each internal node split divides its leaf descendants according to a BetaBinomial distribution (with *β* ≥ 0). In Section 4 we apply Thm. 3.1 to the quartet index and characterize its weak limit. Then, in Section 5 we illustrate the results with simulations. Finally, in the Appendix we provide R code used to simulate from this weak limit.

## 2 Preliminaries

### 2.1 Aldous’s *β*—model for phylogenetic trees

Birth-death models are popular choices for modelling the evolution of phylogenetic trees. However, Aldous (1996, 2001) proposed a different class of models—the so-called *β*–model for binary phylogenetic trees.

The main idea behind this model is to consider a (suitable) family 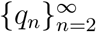 of symmetric, *q_n_*(*i*) = *q_n_*(*n* − *i*), probability distributions on the natural numbers. In particular *q_n_*: {1,…,*n* − 1} → [0,1]. The tree grows in a natural way. The root node of a *n*–leaf tree defines a partition of the *n* nodes into two sets of sizes *i* and *n* − *i* (*i* ∈ {1,…, *n* − 1}). We randomly choose the number of leaves of the left subtree, *L_n_* = *i*, according to the distribution *q_n_* and this induces the number of leaves, *n* − *L_n_*, in the right subtree. We then repeat recursively in the left and right subtrees, i.e. splitting according to the distributions *q_L_n__* and *q_n-L_n__* respectively. Notice that due to *q_n_*’s symmetry the terms left and right do not have any particular meaning attached.

Aldous (1996) proposed to consider a one–parameter, −2 ≤ *β* ≤ ∞, family of probability distributions,

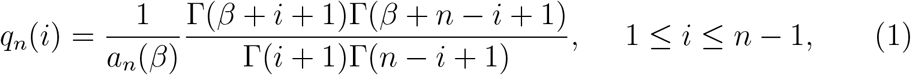

where *a_n_*(*β*) is the normalizing constant and Γ(·) the Gamma function. We may actually recognize this as the BetaBinomial(*n, β* + 1, *β* + 1) distribution and represent

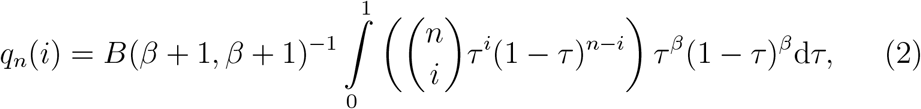

where *B*(*a,b*) is the Beta function with parameters *a* and *b*. Writing informally, from the form of the probability distribution function, Eq. (2), we can see that if we would condition under the integral on *τ*, then we obtain a binomially distributed random variable. This is a key observation that is the intuition for the analysis presented here.

Particular values of *β* correspond to some well known models. The uniform tree model is represented by *β* = −3/2, and the pure birth, Yule, model by *β* = 0.

Of particular importance to our work is the limiting behaviour of the scaled size of the left (and hence right) subtree, *n*^−1^*L_n_*. Aldous (1996) characterizes the asymptotics in his Lemma 3,

#### Lemma 2.1

*(Aldous, 1996, Lemma 3 for β* > −1)

1. *β* = ∞, 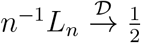;
2. −1 < *β* < ∞, 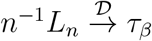, *where τ_β_ has the Beta distribution*

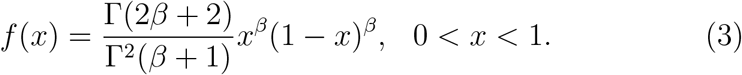

### 2.2 Quartet index

Martínez-Coronado et al. (2018b) introduced a new type of balance index for discrete (i.e. without branch lengths) phylogenetic trees—the quartet index. This index is based on considering the number of so–called quartets of each type made up by the leaves of the tree. A (rooted) quartet is the induced subtree from choosing some four leaves. We should make a point here about the used nomenclature. Usually in the phylogenetic literature a quartet is an unrooted tree on four leaves (e.g. Semple and Steel, 2003). However, here we consider rooted trees and following Martínez-Coronado et al. (2018b) by (rooted) quartet we mean a rooted tree on four leaves. We will from now on write quartet for this, dropping the “rooted” clarification.

For a given tree *T*, let 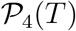 be the set of quartets of the tree. Then, the quartet index of *T* is defined as

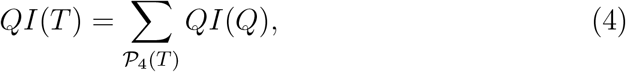

where *QI*(*Q*) assigns a predefined value to a specific quartet. When the tree is a binary one (as here) there are only two possible topologies on four leaves (see Fig. 1). Following Martínez-Coronado et al. (2018b) (their Table 1) we assign the value 0 to *K*_4_ quartets and 1 to *B*_4_ quartets. Therefore, the QI for a binary tree (QIB) will be

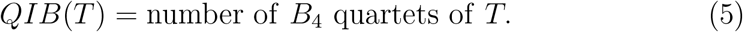

**Figure 1:**
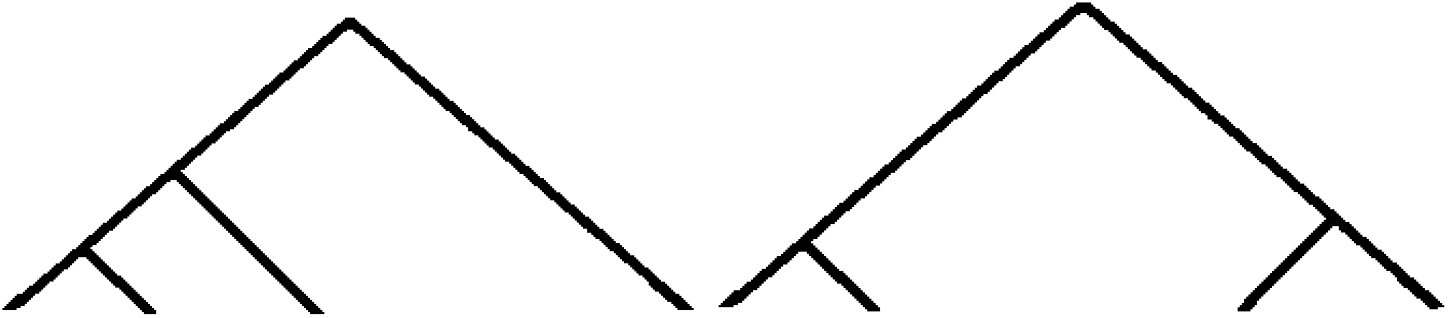
The two possible rooted quartets for a binary tree. Left: *K*_4_ the four leaf rooted caterpillar tree (also known as a comb or pectinate tree), right: *B*_4_ the fully balanced tree on four leaves (also known as a fork, see e.g. Chor and Snir, 2007, for some nomenclature).

Importantly for us Martínez-Coronado et al. (2018b) show in their Lemma 4 that for *n* > 4, the quartet index has a recursive representation as

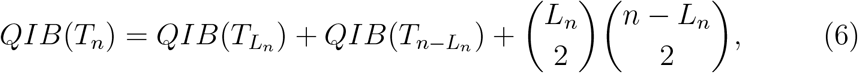

where *T_n_* is the tree on *n* leaves.

Martínez-Coronado et al. (2018b) considered various models of tree growth, Aldous’s *β*–model, Ford’s α–model (Ford, 2005, but see also Martínez-Coronado et al. (2018a)) and Chen–Ford–Winkel’s α–γ–model (Chen et al., 2009). In this work we will focus on the Aldous’s *β* ≥ 0-model of tree growth and characterize the limit distribution, as the number of leaves, *n*, grows to infinity, of the QI. We will take advantage of the recursive representation of Eq. (6) that allows for the usage of the powerful contraction method.

We require knowledge of the mean and variance of the QI and Martínez-Coronado et al. (2018b) show for Aldous’s *β*–model that these are (their Corollaries 4 and 7)

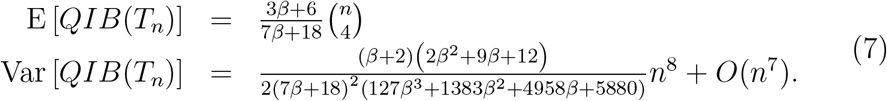

## 3 Contraction method approach

Consider the space *D* of distribution functions with finite second moment and first moment equalling 0. On *D* we define the Wasserstein metric

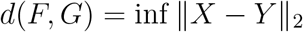

where ||·||_2_ denotes the *L*_2_ norm and the infimum is over all *X* ~ *F, Y* ~ *G*.

Notice that convergence in *d* induces convergence in distribution.

For 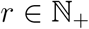 define the transformation *S*: *D* → *D* as

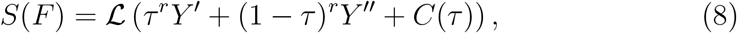

where *Y*′, *Y*″,*τ* are independent, *Y*′, *Y*″ ~ *F, τ* ∈ [0,1] whose distribution is not a Dirac *δ* at 0 nor 1, satisfying, for all *n*,

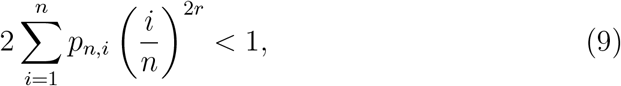

where *p_n,i_* = *P*((*i* − 1)/*n* < *τ* ≤ *i/n*) and the function *C*(·) is of the form

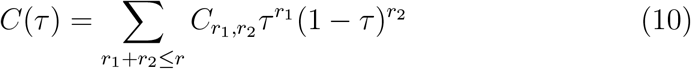

for some constants *C*_*r*_1_,*r*_2__ and furthermore satisfies *E*[*C*(*τ*)] = 0. By Rösler (1992)’s Thms. 3 and 4 *S* is well defined, has a unique fixed point and for any *F* ∈ *D* the sequence *S^n^*(*F*) converges exponentially fast in the *d* metric to *S*’s fixed point. Using the exact arguments to show Rösler (1991)’s Thm. 2.1 one can show that the map *S* is a contraction. Only the Lipschitz constant of convergence will differ being 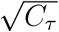, where *C_τ_* = max{E[*τ*^2*r*^], E[(1 − *τ*)^2*r*^]} in our case. Notice that as *τ* ∈ [0,1] and is non-degenerate at the edges, then *C_τ_* < 1 and we have a contraction.

We now state the main result of our work. We show weak convergence, with a characterization of the limit for a class of recursively defined models.

### Theorem 3.1

*(cf. Rösler, 1991, Thm. 3.1) For n* ≥ 2, *β* > 0 *let L_n_* ∈ {1,…, *n* − 1} *be such that* (*L_n_* − 1) *is* BetaBinomial(*n* − 2, *β* + 1, *β* + 1) *distributed and τ* ~ Beta(*β* + 1,*β* + 1) =: *F_τ_ distributed. Starting from the Dirac δ at* 0, *i.e. Y*_1_ = 0 *and convention* BetaBinomial(0,*β* + 1,*β* + 1) = *δ*_0_, *for* 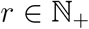 *such that the condition of Eq. (9) is met with the previous choice of F_τ_, define recursively the sequence of random variables*,

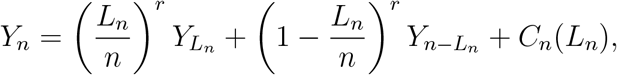

*where the function C_n_*(·) *is of the form*

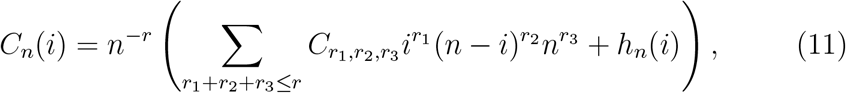

*where* E [*C_n_*(*L_n_*)] = 0 *and* sup_*i*_ *n*^−*r*^*h_n_*(*i*) → 0. *If* 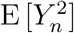 *is uniformly bounded then, the random variable Y_n_ converges in the Wasserstein d–metric to the random variable Y*_∞_ *whose distribution satisfies the unique fixed point of S (Eq. 8)*.

Notice that as *Y*_1_ = 0 and by the definition of the recursion we will have E[*Y_n_*] = 0 for all *n*.

The Yule tree case will be the limit of *β* = 0 and this case the proof of the result will be more straightforward (as commented on in the proof of Thm. 3.1).

Notice that 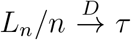. It would be tempting to suspect that Thm. 3.1 should be the conclusion of a general result related to the contraction method (as presented in Eq. (8.12), p. 351 Drmota, 2009). However, to the best of my knowledge, general results assume *L*_2_ convergence of *L_n_/n* (e.g. Thm. 8.6, p. 354 Drmota, 2009), while in our phylogenetic balance index case we will have only convergence in distribution. In such a case it seems that convergence has to be proved case by case (e.g. examples in Rachev and Rüschendorf, 1995). Here we show the convergence of Thm. 3.1 along the lines of Rösler (1991).

We first derive a lemma that controls the non–homogeneous part of the recursion, i.e. *C_n_*(·) as defined in Eq. (11).

### Lemma 3.1

*(cf. Rösier, 1991, Prop. 3.2) Let C_n_*: {1,…,*n* − 1} → ℝ *be as in Eq. (11). Then*

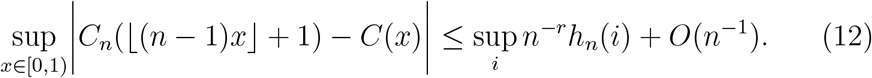

Proof For 1 ≤ ⌊(*n* − 1)*x*⌋ +1 ≤ *n* − 1 and writing *i* = ⌊(*n* − 1)*x*⌋ + 1 we have due to the representation of Eqs. (10) and (11)

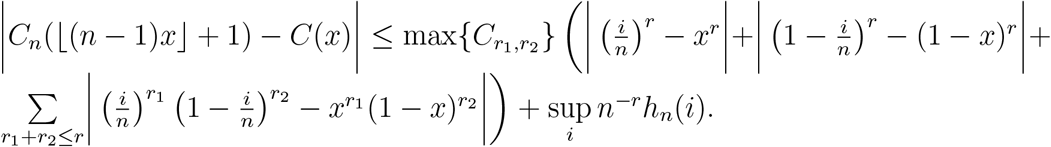

Bounding the individual components, as by construction *x* cannot differ from *i/n* by more than 1/*n*

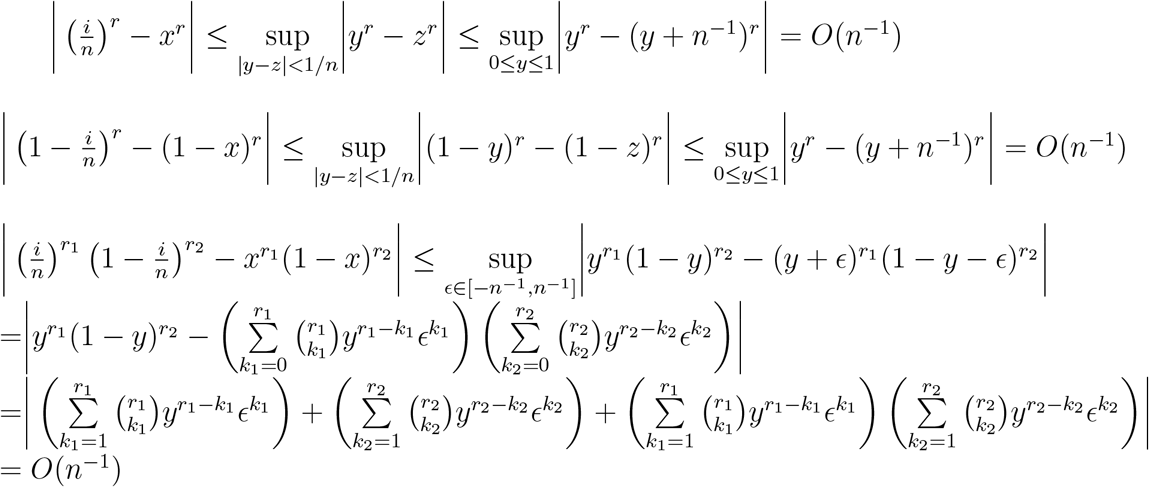

### Lemma 3.2

*(cf. Rösler, 1991, Prop. 3.3) Let a_n_, b_n_, p_n,i_*, 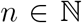 *be three sequences such that* 0 ≤ *b_n_* → 0 *with n*, 0 ≤ *p_n,i_* ≤ 1,

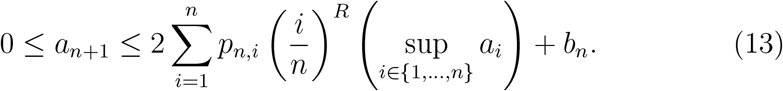

*and*

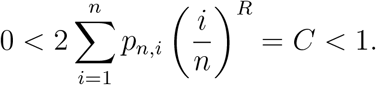

*Then* lim_*n*→∞_ *a_n_* = 0.

Proof The proof is exactly the same as Rösler (1991)’s proof of his Proposition 3.3. In the last step we will have with *a*:= lim sup *a_n_* < ∞ the sandwiching for all *ϵ* > 0

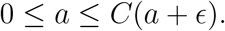

Having Lemmata 3.1 and 3.2 we turn to showing Thm. 3.1.

Proof[Proof of Thm. 3.1] Denote the law of *Y_n_* as 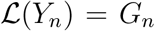. We take *Y*_∞_ and 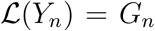 independent and distributed as *G*_∞_, the fixed point of *S*. Then, for *i* = 1,…,*n* − 1 we choose independent versions of *Y_i_* and 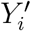. We need to show *d*^2^(*G_n_,G*_∞_) → 0. As the metric is the infimum over all pairs of random variables that have marginal distributions *G_n_* and *G*_∞_ the obvious choice is to take *Y_n_, Y*_∞_ such that *L_n_/n* will be close to *τ* for large *n*. Rösler (1991) was considering the Yule model (*β* = 0) and there *τ* ~ Unif[0,1] and *L_n_* is uniform on {1,…, *n* − 1}. Hence, ⌊(*n* − 1)*τ*⌋ + 1 will be uniform on {1,…,*n* − 1}, remember *P*(*τ* = 1) = 0, and 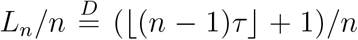. However, when *β* > 0 the situation complicates. For a given *n*, (*L_n_* − 1) is BetaBinomial(*n* − 2,*β* + 1,*β* + 1) distributed (cf. Eq. 1 and Aldous (1996) Eqs. 1 and 3). Hence, if *τ* ~ Beta(*β* + 1,*β* + 1) and (*L_n_* − 1) ~ BetaBinomial(*n* − 2, *β* + 1, *β* + 1) we do not have 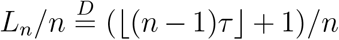 exactly. We may bound the Wasserstein metric by any coupling that retains the marginal distributions of the two random variables. Therefore, from now on we will be considering a version, where conditional on *τ*, the random variable (*L_n_* − 1) is Binomial(*n* − 2, *τ*) distributed. Let *r_n_* be any sequence such that *r_n_/n* → 0 and 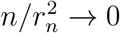, e.g. *r_n_* = *n* ln^−1^ *n*. Then, by Chebyshev’s inequality

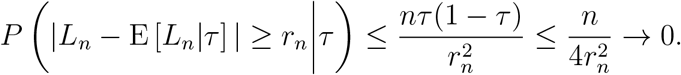

We now want to show *d*^2^(*G_n_, G*_∞_) → 0 and we will exploit the above coupling in the bound

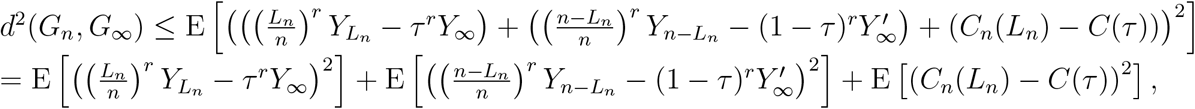

where *Y*_∞_,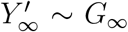 are independent. Remember that E[*Y_i_*] = E[*Y*_∞_] = 0 so that the expectation of the cross products disappears.

Our main step is to have a bound where the *L_n_/n* term is replaced by some transformation of *τ*. Let 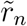 be a (appropriate) random integer in {±1,…, ±⌈*r_n_*⌉} and we may write (with the chosen coupling of *L_n_* and *τ*),

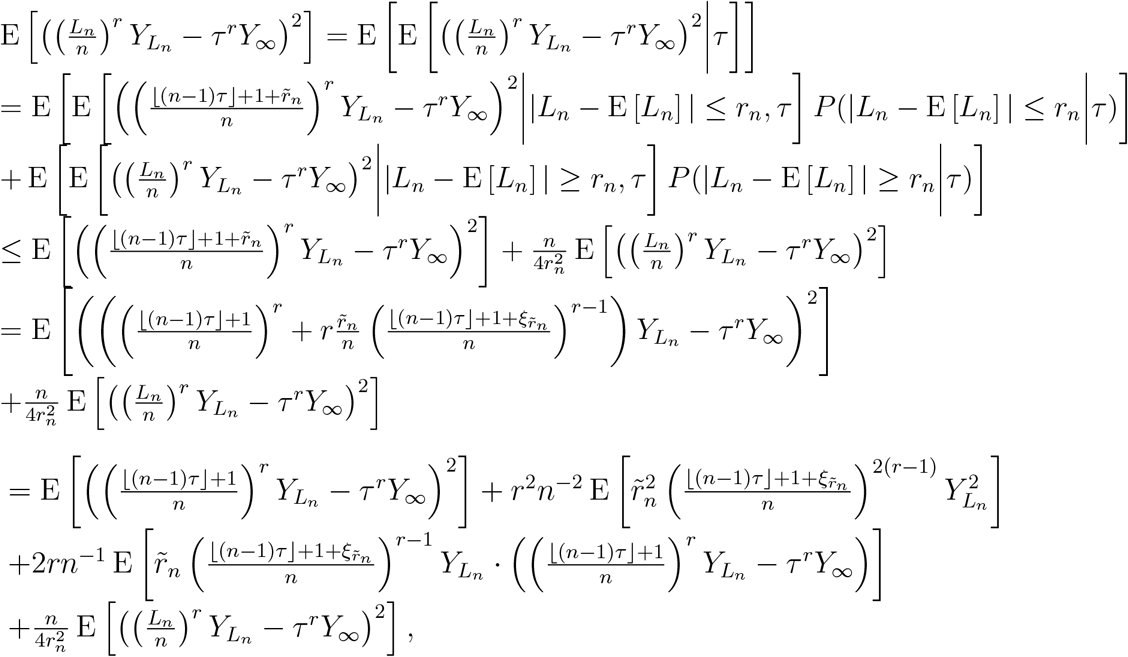

where 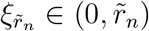 is (a random variable) such that the mean value theorem holds (for the function (·)^*r*^). As *Y_n_, Y*_∞_ have uniformly bounded second moments, 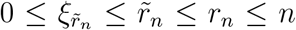, by assumption *r_n_/n* → 0 and 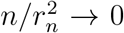, we have that

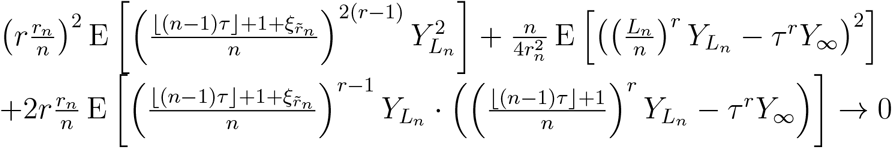

and hence for some sequence *u_n_* → 0 we have,

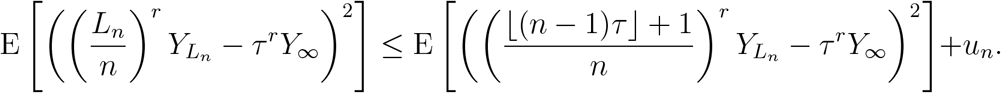

Remembering the assumption sup_*i*_ *n*^−*r*^*h_n_*(*i*) → 0, the other component can be treated in the same way as 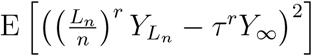 with conditioning on *τ* and then controlling by *r_n_* and Chebyshev’s inequality *L_n_*’s deviation from its expected value. We therefore have for some sequence *v_n_* → 0

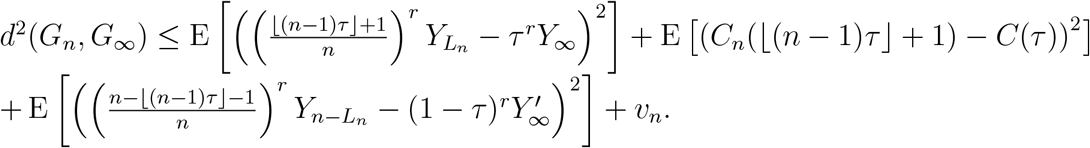

Consider the first term of the right-hand side of the inequality and denote 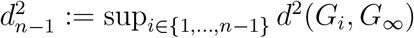

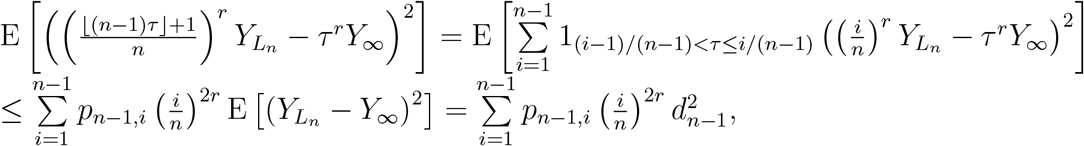

where *p_n,i_* = *P*((*i* − 1)/(*n* − 1) < *τ* ≤ *i*/(*n* − 1)). Invoking Lemmata 3.1, 3.2 and using the assumption of Eq. (9) with *R* = 2*r* we have

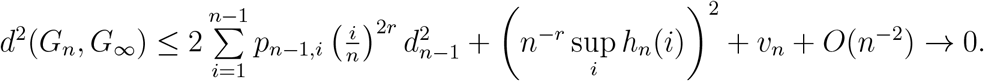

## 4 Limit distribution of the quartet index for Aldous’s *β* ≥ 0–model trees

We show here that the QIB of Aldous’s *β* ≥ 0-model trees satisfies the conditions of Thm. 3.1 with *r* = 4 and hence the QIB has a well characterized limit distribution. We define a centred and scaled version of the QIB for Aldous’s *β* ≥ 0-model tree on *n* ≥ 4 leaves

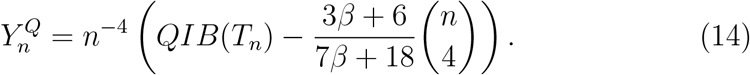

We now specialize Thm. 3.1 to the QIB case and assume *Y*_1_ = *Y*_2_ = *Y*_3_ = 0 for completeness

### Theorem 4.1

*The sequence of random variables* 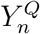 *random variable for trees generated by Aldous’s β-model with β* ≥ 0 *converges with n* → ∞ *in the Wasserstein d-metric (and hence in distribution) to a random variable* 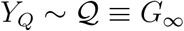 *satisfying the following equality in distribution*

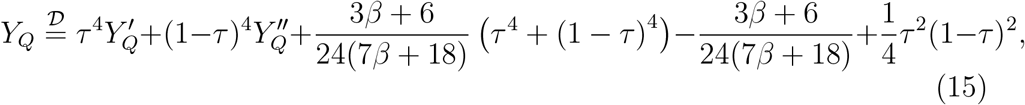

*where τ* ~ *F_τ_ is distributed as the Beta distribution of Eq. (3)*, 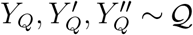 *and* 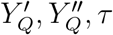 *are all independent*.

Proof Denote by *P*_3_(*x,y*) a polynomial of degree at most three in terms of the variables *x, y*. From the recursive representation of Eq. (6) for *n* > 4

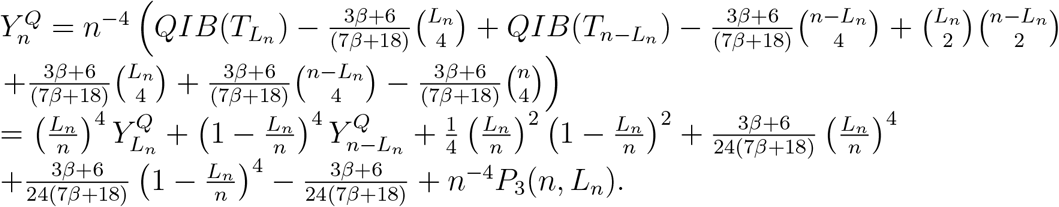

We therefore have *r* = 4 and

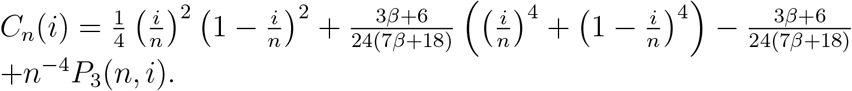

By the scaling and centring we know that 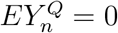 and 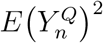 is uniformly bounded by Eq. (7). For the Beta law of *τ* we need to examine for all *i*

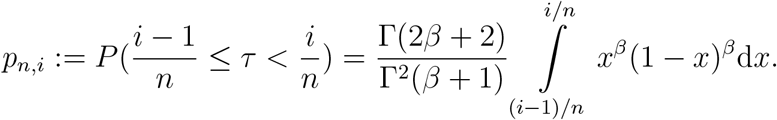

We consider two cases

1. *β* > 0, we have to check if the condition of Eq. (9) is satisfied. Let

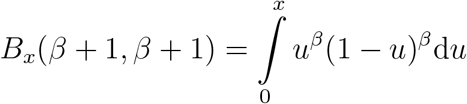

be the incomplete Beta function. Then,

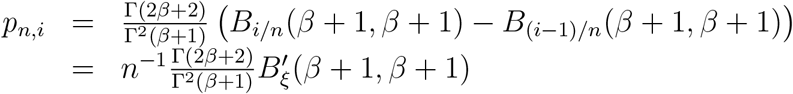

by the mean value theorem for some *ξ* ∈ ((*i* − 1)/*n, i/n*). Obviously

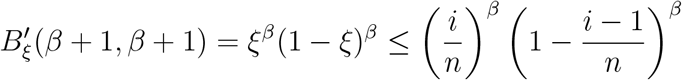

and now

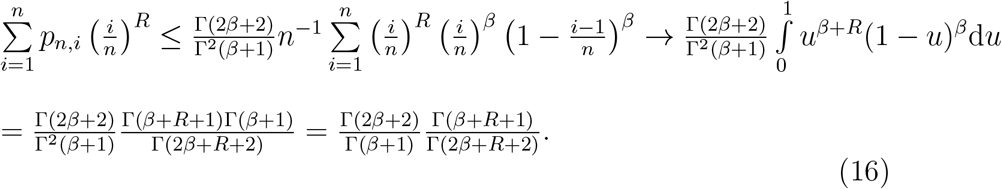 Take 1 < *R*_1_ < *R*_2_ and consider the ratio

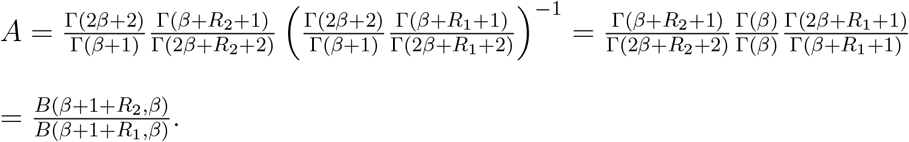 The ratio *A* < 1 as the Beta function is decreasing in its arguments— hence the derived upper bound in Eq. (16) is decreasing in *R*. For *R* =1 the bound equals

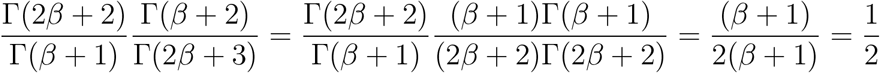

and hence for all *R* > 1 and all *β* > 0

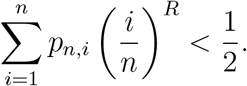 As in our case we have *r* ≥ 1, then for *R* = 2*r* ≥ 2 the assumptions of Lemma 3.2 are satisfied and the statement of the theorem follows through.
2. *β* = 0, then directly *p_n,i_* = *n*^−1^, Eq. (9) and assumptions of Lemma 3.2 are immediately satisfied and the statement of the theorem follows through. This is the Yule model case, in which the proof of the counterpart of Thm. 3.1 is much more straightforward, as mentioned before.

### Remark 4.1

*When β* < 0 *the process L_n_/n seems to have a more involved asymptotic behaviour (cf. Lemma 3 of Aldous, 1996, in β* ≤ −1 *case). Furthermore, the bounds applied here do not hold for β* < 0. *Therefore, this family of tree models (including the important uniform model, β = −3/2) deserves a separate study with respect to its quartet index*.

## 5 Comparing with simulations

To verify the results we compared the simulated values from the limiting theoretical distribution of *Y_Q_* with scaled and centred values of Yule tree QI values. The 500-leaf Yule trees were simulated using the `rtreeshape()` function of the apTreeshape R package and Tomas Martínez–Coronado’s inhouse Python code. Then, for each tree the QI value was calculated by Gabriel Valiente’s and Tomas Martínez–Coronado’s in–house programs. The raw values QIB(Yule_500_) were scaled and centred as

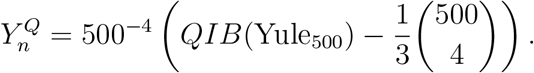

The *Y_Q_* values were simulated using the proposed by Bartoszek (2018) heuristic Algorithm 3 (R code in Appendix). The results of the simulation are presented in Fig. 2.

**Figure 2:**
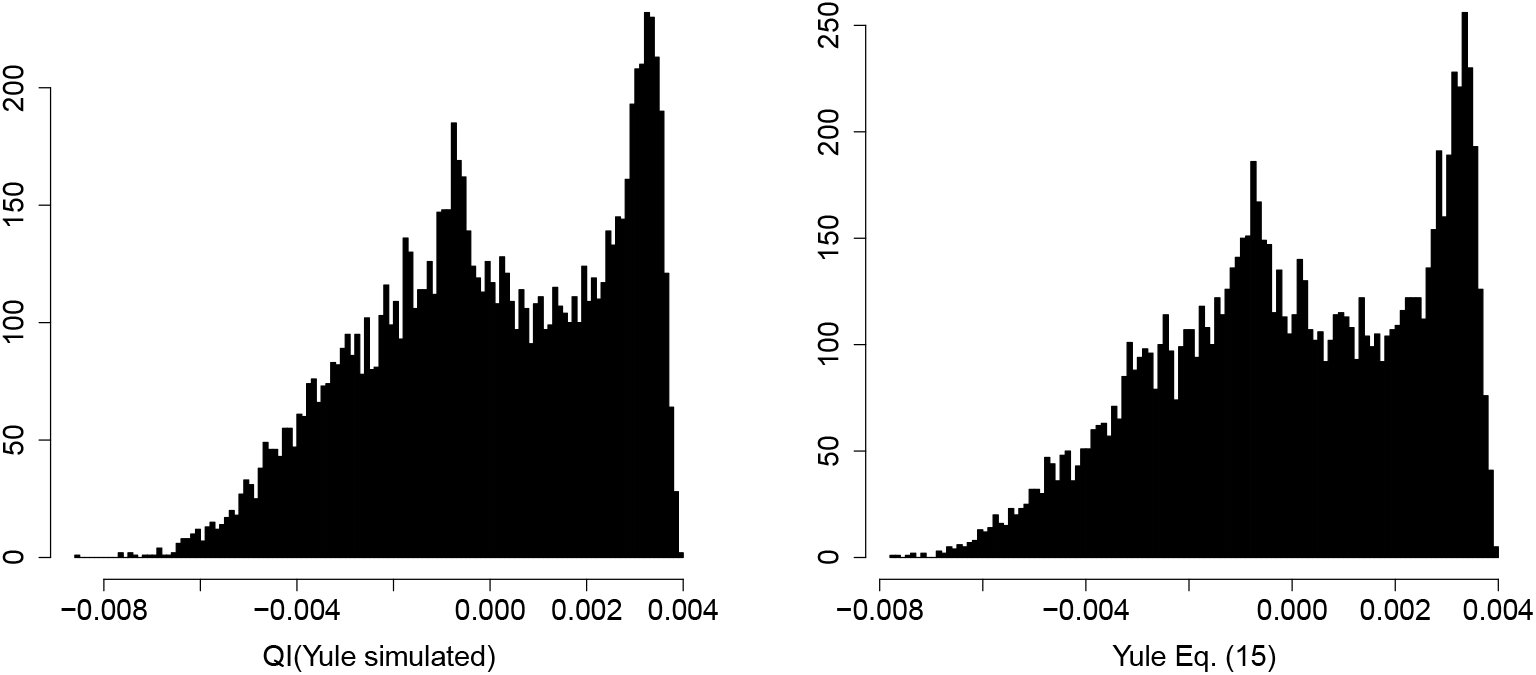
Left: histogram of scaled and centred simulated values of the QIB for the Yule tree, 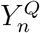, right: histogram of *Y_Q_* for the Yule model, *β* = 0. The mean, variance, skewness and excess kurtosis of the simulated values are −3.177 · 10^−6^, 6.321 · 10^−6^, −0.308, −0.852 (left, simulated values) and 1.682·10^−5^, 6.38· 10^−6^, −0.317, −0.834 (right, theoretical values by Bartoszek (2018)’s heuristic Algorithm 2 with recursion depth 15). For *β* = 0 the leading constant of the variance in Eq. (7) is 5/(24 · 33075) ≈ 6.299 · 10^−6^.

## Acknowledgments

I would like to thank the whole Computational Biology and Bioinformatics Research Group of the Balearic Islands University for hosting me on multiple occasions, introducing me to problems related with the quartet index and for valuable comments on this manuscript. The simulated values of the quartet index for the Yule tree were provided by Gabriel Valiente and Tomás Martínez–Coronado. I was supported by the Knut and Alice Wallenberg Foundation and am by the Swedish Research Council (Veten-skapsradet) grant no. 2017–04951. My visits to the Balearic Islands University were partially supported by the the G S Magnuson Foundation of the Royal Swedish Academy of Sciences (grants no. MG2015–0055, MG2017–0066) and The Foundation for Scientific Research and Education in Mathematics (SVeFUM).

## Appendix: R code for simulating from the limit distribution of the quartet index

~~~
fCtau_QIB<−function (x, beta=0){
    PB4beta<−(3*beta + 6)/ (7* beta + 18);
    PB4beta * (x^4) / 2 4+PB4beta * ((1 − x)^4)/24 − PB4beta / 24 + 0.25 * x * x *(1 − x)^2
}
fdistribution_limitQIB <−function (num. iter=10,popsize = 10000, Y0=0){
    replicate (popsize, fdraw_limitQIB (num. iter, Y0))
}
fdraw_limitQIB<−function (num. iter = 15,Y0=0){
    res <−0
    if (num. iter ==0){
        Y1<−Y0
        Y2<−Y0
    }
    else {
           Y1<− fdraw_limit QIB (num. iter −1, Y0)
           Y2<− fdraw_limit QIB (num. iter −1, Y0)
    }
    tau<− runif(1)
    res<−((tau)^4) * Y1 + ((1 − tau)^4) * Y2+fCtau_QIB (tau)
 res
}
popsize < −10000 ## size of sample for histogram
num. iter <−15 ## depth of the recursion
Y0 <− 0 ## initial value
vlimitQIB<− fdistribution_limitQIB (num. iter=num. iter, popsize=popsize)
~~~

